# A cell-based, spike protein binding assay highlights differences in antibody neutralising capacity for SARS-CoV-2 variants

**DOI:** 10.1101/2022.06.24.496409

**Authors:** Neale Harrison, Lauren Richardson, Chiara Pallini, Ines Morano, Elizabeth Jinks, Jamie Cowley, Hujo Chan, Harriet J Hill, Cristina Matas de las Heras, Ana Teodosio, Andrea S Lavado, Timothy R Dafforn, Dimitris K Grammatopoulos, John Gordon, Catherine A Brady, Lawrence S Young, Nicholas M Barnes, Zania Stamataki, Omar S Qureshi

## Abstract

The engagement of the SARS-CoV-2 spike protein with ACE2 is a critical step for viral entry to human cells and accordingly blocking this interaction is a major determinant of the efficacy of monoclonal antibody therapeutics and vaccine-elicited serum antibodies. The emergence of SARS-CoV-2 variants necessitates the development of adaptable assays that can be applied to assess the effectiveness of therapeutics. Through testing of a range of recombinant spike proteins, we have developed a cell based, ACE2/spike protein binding assay that characterises monoclonal anti-spike protein antibodies and neutralising antibodies in donor serum. The assay uses high-content imaging to quantify cell bound spike protein fluorescence. Using spike proteins from the original ‘Wuhan’ SARS-CoV-2 virus, as well as the delta and omicron variants, we identify differential blocking activity of three monoclonal antibodies directed against the spike receptor binding domain. Importantly, biological activity in the spike binding assay translated to efficacy in a SARS-CoV-2 infection assay. Hence, the spike binding assay has utility to monitor anti-spike antibodies against the major known SARS-CoV-2 variants and is readily adaptable to quantify impact of antibodies against new and emerging SARS-CoV-2 variants.

## Introduction

The on-going SARS-CoV-2 pandemic necessitates the development of tools that can support the advance of therapeutics that are efficacious against both current and future viral variants (1). Quantifying the activity of neutralising antibodies, either recombinant monoclonals or those elicited by vaccination, is therefore of considerable importance for drug development and for monitoring effective immunity (2,3). Emerging SARS-CoV-2 variants may exhibit escape or reduced neutralisation by current monoclonal antibodies or from those antibodies generated by vaccination generated to target earlier viral strains (4) and knowledge of these limitations informs health and political strategy in the face of societal challenges arising from the pandemic.

The SARS-CoV-2 spike protein engagement with the host ACE2 receptor is a key step in viral entry (5) and is therefore a major target for therapeutic interventions (6,7,8,9) as well as a relevant mechanism to study efficacy of neutralising antibodies in serum (10, 11). In addition, spike protein priming requires the serine protease TMPRSS2 (5). Functional virus neutralisation experiments require biosafety level 3 facilities in addition to the requirement for propagation of strains prior to infection studies, which carry inherent risk. Reduced risk procedures using pseudovirus still requires packaging and expression of viral particles (12). We therefore sought to develop a reductionist cell-based model, suitable for high-throughput screening, that quantifies inhibition of SARS-CoV-2 spike protein binding to mammalian cells suitable to assist therapeutic development and immune monitoring.

## Results and Discussion

### Binding of SARS-CoV-2 spike protein to cells over-expressing ACE2

Initial experiments investigated the binding of a recombinant His-tagged SARS-CoV-2 spike protein (Spike A) to lung epithelial A549 cell lines, either wild-type (A549-WT), over-expressing ACE2 (A549-ACE2), or over-expressing ACE2 and TMPRSS2 (A549-ACE2-TMPRSS2). The His-tagged spike protein was detected using a mouse IgG anti-His antibody, followed by an AF488-conjugated anti-mouse IgG secondary antibody, followed by visualisation by high-content imaging (**Figure 1A**) quantifying the intensity of spike protein labelling per cell **(Figure 1C**). Whilst labelling of wild-type A549 cells with spike protein was close to background levels, clear punctate labelling of spike protein was evident on both A549 cells over-expressing ACE2, and on those over-expressing ACE2 and TMPRSS2. Spike protein labelling was lower on A549-ACE2-TMPRSS2 cells compared to A549-ACE2 and may simply reflect differences in ACE2 expression between the cell lines. Subsequent use of a spike protein within a detergent micelle (Spike F), which likely contains the trimeric spike in its native configuration, exhibited clear labelling as detected by an anti-RhoD1A4 antibody and an AF488-conjugated anti-mouse IgG secondary antibody (**Figure 1B and C**).

**Figure 1.**
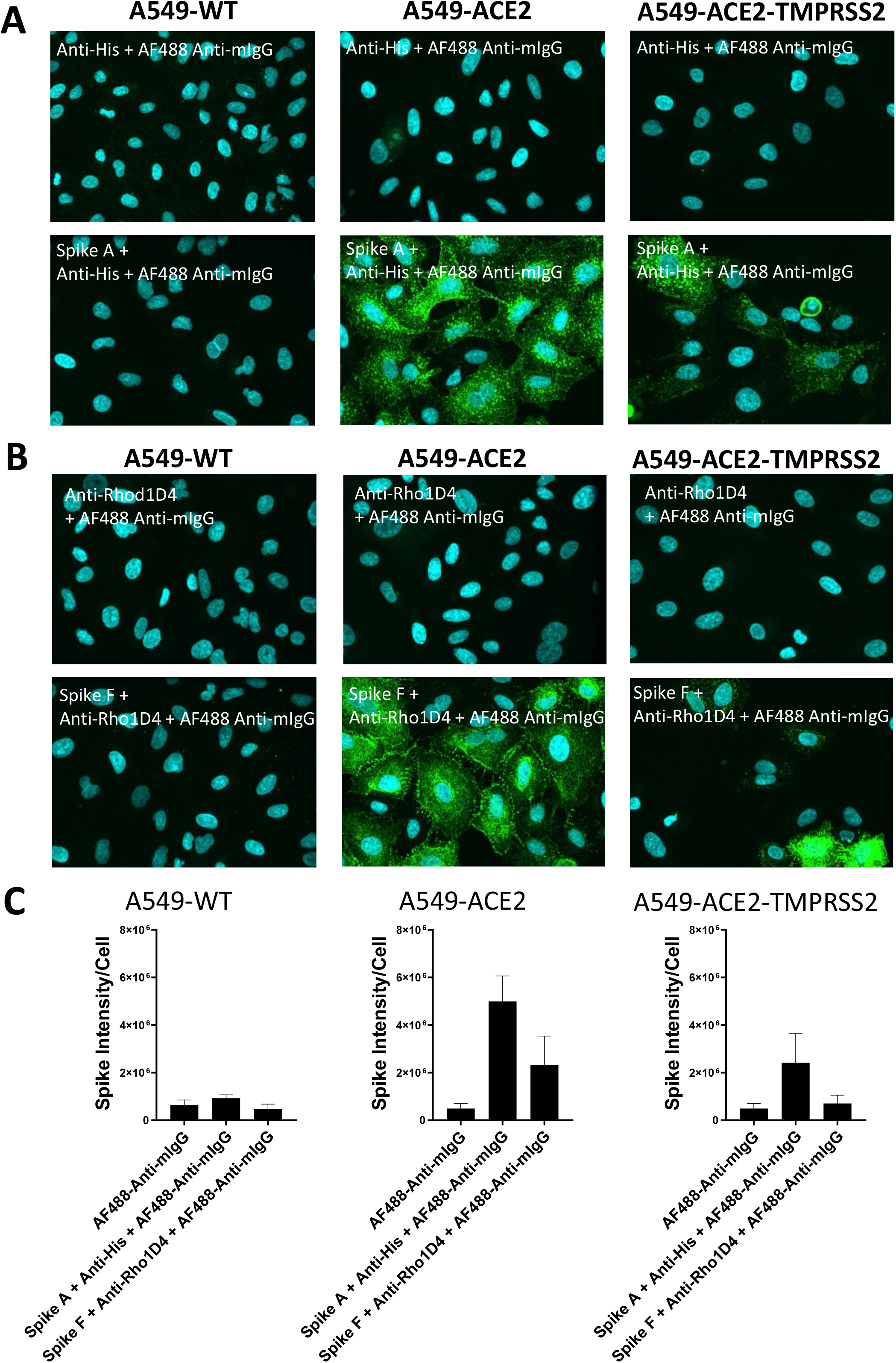
Labelling of ACE2 expressing A549 cells by recombinant spike proteins. (**A**) Representative confocal images showing labelling of wild type (A549-WT), ACE2-overexpressing (A549-ACE2) and ACE2 and TMPRSS2-overexpressing (A549-ACE2-TMPRSS2) A549 cells by recombinant spike protein A (‘Spike A’) and anti-His + AF488-conjugated anti-mIgG detection antibodies. (**B**) Representative confocal images showing labelling of A549 cells (A549-WT), ACE2-overexpressing (A549-ACE2) and ACE2 and TMPRSS2-overexpressing (A549-ACE2-TMPRSS2) A549 cells by recombinant spike protein F (‘Spike F’) and anti-His + AF488-conjugated anti-mIgG detection antibodies (green; nuclei in cyan). (**C**) Quantification of spike protein labelling intensity from images acquired by high content confocal microscopy for cells labelled in (**A**) and (**B**). Data expressed as mean + SEM from three independent experiments for A549-ACE2 and from two independent experiments for A549-WT.

Subsequent evaluation studied a range of recombinant spike proteins (listed in Table 1) to A549-WT and A549-ACE2 cells. Binding to A549-WT was low generally (**Figure 2A**). However, with A549-ACE2 cells, a range of labelling intensities was evident with some recombinant spike proteins exhibiting minimal signal **(Figure 2B)**. These data confirmed that certain recombinant spike proteins allowed clear labelling of A549-ACE2 cells.

**Table 1.** Recombinant spike proteins.

|  | Name | Lineage | Introduced mutations | Supplier | Catalogue number |
| --- | --- | --- | --- | --- | --- |
| **Spike A** | SARS-CoV-2 S protein, His Tag, Super stable trimer | Accession #: QHD43416.1 | F817P, A892P, A899P, A942P, K986P and V987P; R683A and R685A | Acro Biosystems | SPN C52H8 |
| **Spike B** | SARS-CoV-2 S protein (D614G), His,Avitag™, Super stable trimer, biotinylated | Accession #: QHD43416.1 (D614G) | F817P, A892P, A899P, A942P, K986P and V987P; R683A and R685A | Acro Biosystems | SPN-C82E3 |
| **Spike C** | SARS-CoV-2 S protein, His,Avitag™, Super stable trimer, biotinylated | Accession # QHD43416.1) | F817P, A892P, A899P, A942P, K986P and V987P; R683A and R685A | Acro Biosystems | SPN-C82E9 |
| **Spike D** | Spike Sheep Fc-tag (HEK293) | Accession Number: YP_009724390.1 |  | Native antigen company | REC31807 |
| **Spike E** | SARS-CoV-2 S protein, His Tag, Super stable trimer | Accession number: QHD43416.1 | F817P, A892P, A899P, A942P, K986P and V987P; R683A and R685A | Acro Biosystems | SPN-C52H9 |
| **Spike F** | SARS CoV-2-full-length Spike B.1.1.7 | Alpha; B.1.1.7 | furin cleavage site "RRAR" mutated to "GSAG"; KV986PP | Cube Biotech | 28716 |
| **Spike G** | SARS-CoV-2 (2019-nCoV) Spike RBD(N501Y)-His Recombinant Protein | Accession number: YP_009724390.1 |  | Sino Biological | 40592-V08H82-B |
| **Spike H** | SARS-CoV-2 Spike His Protein, CF | Accession number: YP_009724390.1 | Arg682Ser, Arg685Ser, Lys986Pro and Val987Pro | R&D systems | 10549-CV |
| **Spike I** | SARS-CoV-2 Spike (GCN4-IZ) His Protein, CF | Accession number: YP_009825051.1 | K968P, V969P | R&D systems | 10581-CV |
| **Spike J** | SARS-CoV-2 Spike (GCN4-IZ) His Protein, CF | Accession number: YP_009724390.1 | Arg682Ser, Arg685Ser, Lys986Pro and Val987Pro | R&D systems | 10638-CV |
| **Spike K** | SARS-CoV-2 Spike (D614G) His Protein, CF | Accession number: YP_009724390.1 | Asp614Gly, Arg682Ser, Arg685Ser, Lys986Pro, Val987Pro | R&D systems | 10620-CV |
| **Spike L** | SARS-CoV-2 Spike Trimer (Delta) | Delta variant (Pango lineage: B.1.617.2; Accession number: | F817P, A892P, A899P, A942P, K986P, V987P; R683A and R685A | Acro Biosystems | SPN-C52He |
| **Spike M** | SARS-CoV-2 Spike RBD | Omicron variant (Pango lineage B.1.1.529); |  | Acro Biosystems | SPD-C522e |

**Figure 2.**
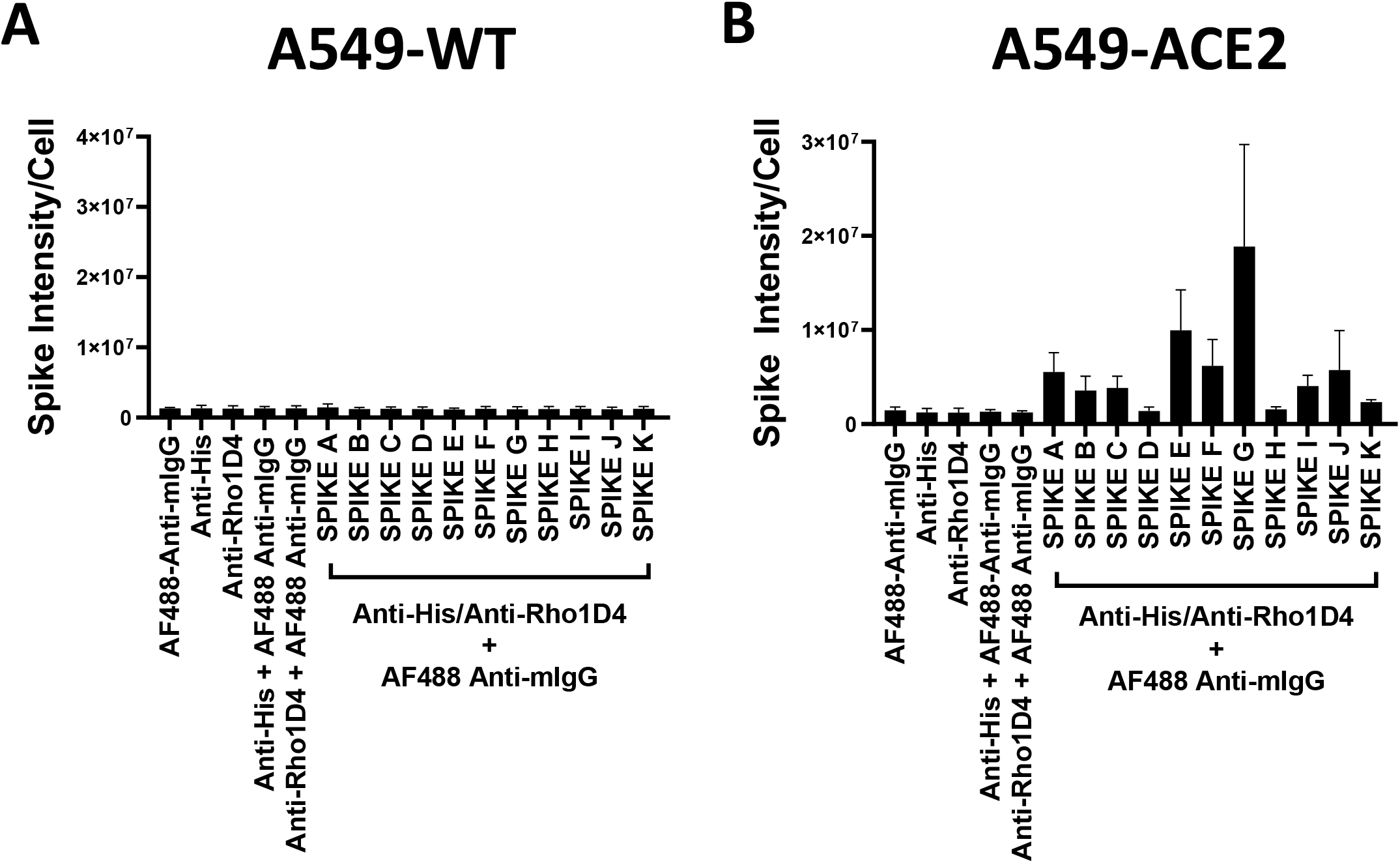
Comparison of recombinant spike proteins. Labelling of A549-WT (**A**) and A549-ACE2 (**B**) cells by recombinant spike proteins (listed in Table 1). Spike proteins were detected using anti-His + AF488-conjugated anti-mouse IgG antibodies except for Spike F which was detected using anti-Rho1D4 + AF488-conjugated anti-mouse IgG antibodies, followed by visualisation and quantification of spike protein labelling intensity by high content confocal microscopy. Data expressed as mean + SEM from three independent experiments.

### Blockade of spike protein binding by monoclonal antibodies

To test the utility of this assay for monoclonal antibody screening, the impact of a range of commercially available anti-spike antibodies (Table 2) on labelling of ACE2-A549 cells by Spike A **(Figure 3A)** and Spike F **(Figure 3B)** were investigated. Antibodies RBD1, RBD5, RBD7 and RBD8, consistently reduced labelling of both Spike A and Spike F, and three of these antibodies were selected for further study. Titration of RBD1, RBD7 and RBD8 revealed concentration-dependent blockade of spike protein A (**Figure 4A)** and E (**Figure 4B**) labelling allowing quantification of IC_50_ values for each of the antibodies (**Figure 4 and Table 3**). To assess the translational relevance of the assay, the ability of RBD1, RBD7 and RBD8 to neutralise SARS-CoV-2 infection of Vero cells was investigated. Similar to the spike labelling results with Spike A (**Figure 4A**), RBD1 exhibited the lowest potency to neutralise viral infection whereas both RBD7 and RBD8 effectively prevented viral infection, with RBD8 exhibiting greater potency than in the spike binding assay (**Figure 5 and Table 4**).

**Table 2.** Recombinant Antibodies.

|  | Name | Supplier | Catalogue Number |
| --- | --- | --- | --- |
| RBD1 | SARS-CoV1/2 Spike RBD Llamabody antibody | R&D Systems | LMAB10541 |
| RBD2 | SARS-CoV2 Spike protein (RBD) recombinant human monoclonal antibody | Thermofisher Scientific | T01KHu |
| RBD3 | SARS-CoV2 Spike protein (RBD) polyclonal antibody | Thermofisher Scientific | PA5-116915 |
| RBD4 | SARs-CoV-2 Spike Protein (S-ECD/RBD) Monoclonal antibody (bcb01) | Thermofisher Scientific | MA5-35948 |
| RBD5 | SARs-CoV-2 Spike Protein (S-ECD/RBD) Monoclonal antibody (bcb02) | Thermofisher Scientific | MA5-35949 |
| RBD6 | SARs-CoV-2 Spike Protein (S-ECD/RBD) Monoclonal antibody (bcb03) | Thermofisher Scientific | MA5-35950 |
| RBD7 | Recombinant Anti-SARS-CoV2 Spike RBD antibody (CV30) | Abcam | ab277513 |
| RBD8 | Recombinant Anti-SARS-CoV2 Spike RBD antibody (HL1003) | Abcam | ab281303 |
| RBD10 | Anti-spike protein IgG1 Fc silent (CV1) | Absolute Antibody | Ab02018-10.3 |
| RBD11 | Anti-spike protein IgG1 (CV1) | Absolute Antibody | AB02018-10.0 |
| RBD12 | Anti-spike protein IgG1 (CR3022) | Absolute Antibody | AB01680-10.0 |
| RBD13 | Anti-spike protein IgG1 Fc silent (CR3022) | Absolute Antibody | AB01680-10.3 |
| RBD14 | SARS-CoV1/2 Spike RBD Llamabody antibody | R&D Systems | LMAB10869 |

**Table 3.** IC50 values ± SEM from four independent experiments; RBD1, RBD7 and RBD8; Spike A and E.

|  | IC<sub>50</sub> |
| --- | --- |
| RBD1; Spike A | 2.65 ± 1.63 |
| RBD7; Spike A | 1.03 ± 0.35 |
| RBD8; Spike A | 1.53 ± 0.52 |
| RBD1; Spike E | n.d. |
| RBD7; Spike E | 0.75 ± 0.14 |
| RBD8; Spike E | 1.75 ± 0.50 |

**Table 4.** IC50 values; SARS-CoV-2 infection of Vero cells.

|  | IC<sub>50</sub> |
| --- | --- |
| RBD1 | 0.332 |
| RBD7 | 0.133 |
| RBD8 | 0.003 |

**Figure 3.**
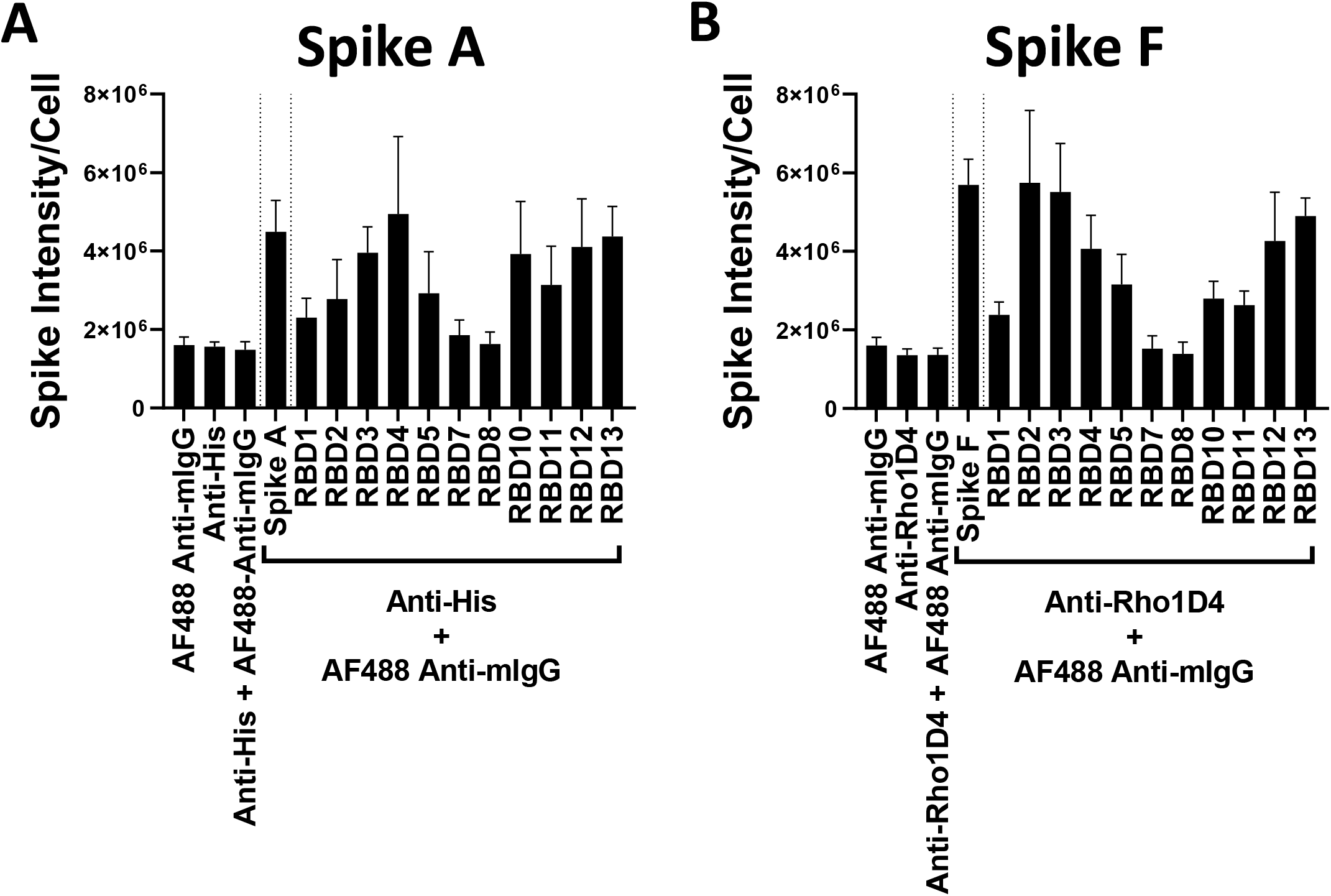
Antibody blockade of spike protein binding. Labelling of A549-ACE2 cells by recombinant spike protein A **(A)** or F **(B)** that had been pre-incubated in the absence or presence of anti-spike antibodies (as listed in table 2) at 10 μg/mL, followed by detection using anti-His or anti-Rho1D4 + AF488-conjugated anti-mouse IgG antibodies, and visualisation and quantification of spike protein labelling intensity by high content confocal microscopy. Data expressed as mean + SEM from three independent experiments.

**Figure 4.**
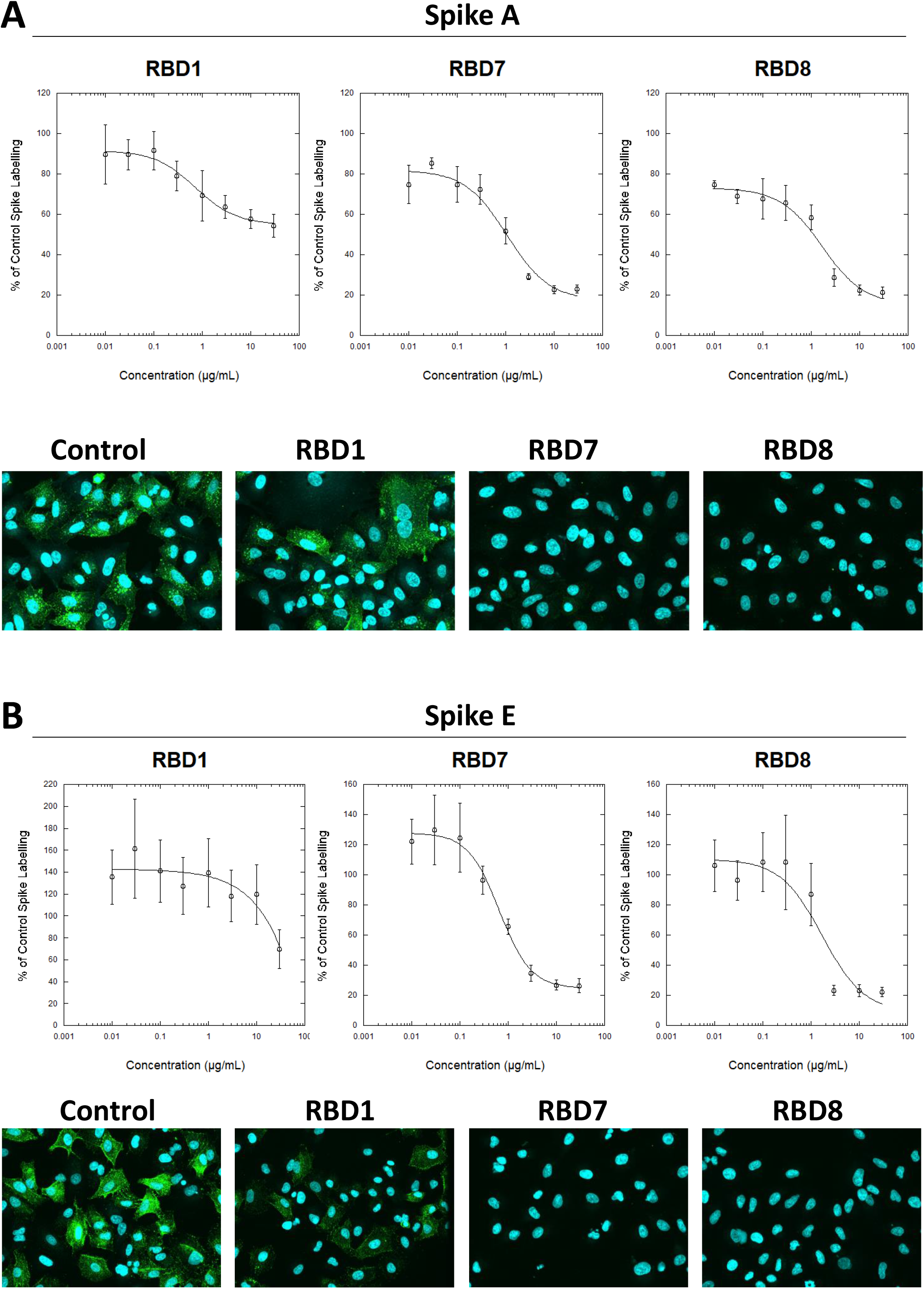
Characterisation of spike blocking antibodies. Labelling of A549-ACE2 cells by recombinant spike protein A (**A**) or E **(B**) that had been pre-incubated in the absence or presence of the indicated concentrations of antibodies RBD1, RBD7 or RBD8, followed by detection using anti-His + AF488-conjugated anti-mouse IgG antibodies, and visualisation and quantification of spike protein labelling intensity by confocal microscopy. Plots show data expressed as mean ± SEM of three independent experiments. Solid line represents nonlinear regression using a 4-parameter logistic equation. Representative confocal images show labelling of cells under control conditions or with spike protein pre-incubated with 10 μg/mL RBD1, RBD7 or RBD8.

**Figure 5.**
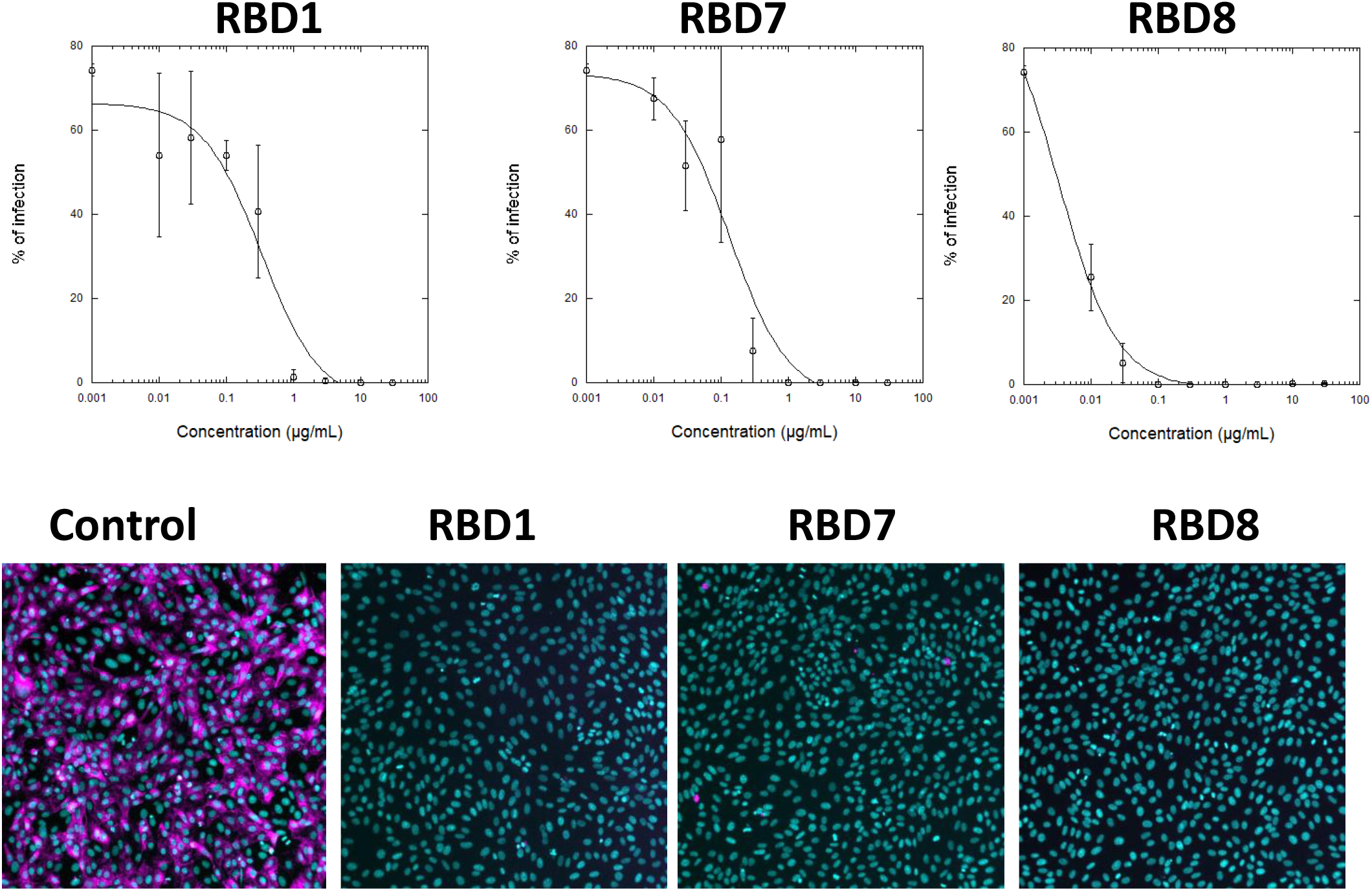
Impact of spike blocking antibodies on SARS-CoV-2 infection of Vero cells. Vero cells were infected with 3.3×10^3^ IU/ml of hCOV-19/England/2/2020 virus isolate in the absence or presence of the indicated concentrations of RBD1, RBD7 or RBD8. Infection rates were assessed at 24 h by staining Vero cells for viral spike protein (magenta) and counterstaining nuclei with Hoechst (cyan). Plots show data expressed as mean ± SD of replicate wells. Solid line represents nonlinear regression using a 4-parameter logistic equation. Representative confocal images show labelling of cells under control conditions or with spike protein pre-incubated with 10 μg/mL RBD1, RBD7 or RBD8.

To expand the potential utility of the spike binding assay, performance with plasma from individuals was evaluated with anti-spike IgG/A/M determined by ELISA (**Figure 6A**). A value ≥ 1.0 is considered positive in this assay and the plasma samples containing levels of anti-spike IgG/A/M >3 exhibited concentration-dependent spike blocking activity, whereas those with levels <3 did not (**Figure 6B**).

**Figure 6.**
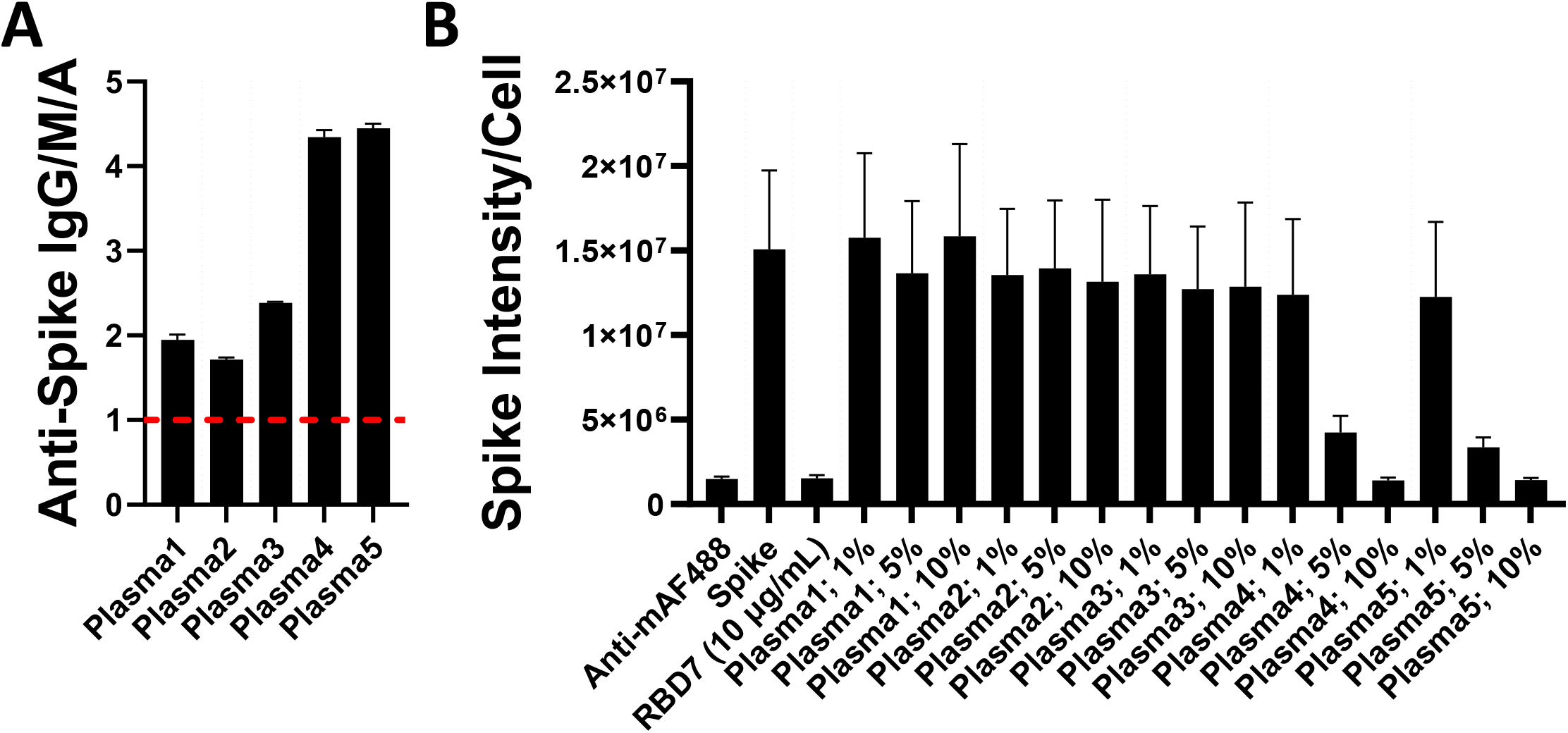
Impact of plasma on spike protein binding. (**A**) Plasma levels of anti-Spike IgG/M/A quantified by ELISA. Data expressed are mean + SD from three replicate wells. Dashed red line indicates threshold for positive for SARS-CoV-2 antibodies. (**B**) Labelling of A549-ACE2 by Spike E pre-incubated in the absence or presence of plasma at 1, 5 or 10%, or RBD7 (10 μg/mL) followed by detection using anti-His + AF488-conjugated anti-mouse IgG antibodies, and visualisation and quantification of spike protein labelling intensity by confocal microscopy. Data expressed as mean + SEM from three independent experiments.

Taken together, these data suggest that the spike binding assay is suitable for both monoclonal antibody characterisation and for monitoring neutralising antibody activity in plasma.

### Blockade of viral variant spike protein binding by monoclonal antibodies

During this study, the emergence of the SARS-CoV-2 variants ‘delta’ and ‘omicron’ highlighted the need to rapidly assess activity of potential therapeutics against the variant spike protein. The activity of RBD7, RBD8 and RBD14 was compared against the delta and omicron spike protein labelling of A549-ACE2 cells. Whilst RBD7, RBD8 and RBD14 effectively reduced labelling by the delta spike protein (**Figure 7A**; **Table 3;** with IC_50_ values of 0.24, 0.76 and 1.47 μg/mL, respectively), only RBD14 displayed a comparable efficacy against the Omicron spike protein (**Figure 7B**; **Table 5;** IC_50_ of 0.97 μg/mL).

**Table 5.** IC50 values ± SEM from three independent experiments; RBD1, RBD7 and RBD8; Spike A and E.

|  | IC<sub>50</sub> |
| --- | --- |
| RBD14; Delta | 0.26 ± 0.1 |
| RBD7; Delta | 0.65 ± 0.30 |
| RBD8; Delta | 1.56 ± 0.36 |
| RBD14; Omicron | 0.95 ± 0.43 |
| RBD7; Omicron | n.d. |
| RBD8; Omicron | 5.52 ± 4.28 |

**Figure 7.**
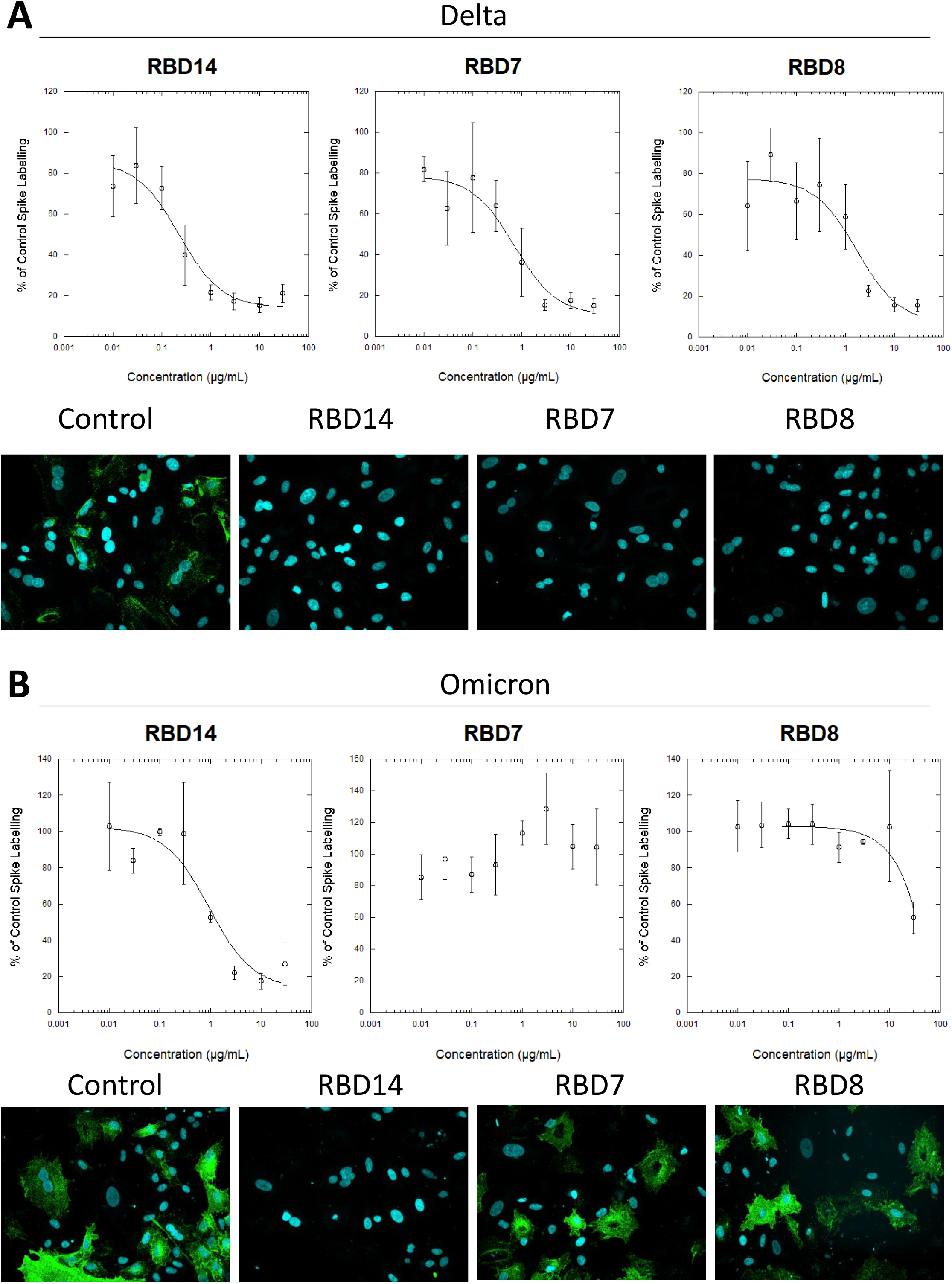
Characterisation of spike blocking antibodies against variants delta and omicron. Labelling of A549-ACE2 cells by recombinant delta spike protein (**A;** ‘Spike L’) and omicron spike protein (**B;** ‘Spike M’) pre-incubated in the absence or presence of the indicated concentrations of antibodies RBD1, RBD7 or RBD8, followed by detection using anti-His + AF488-conjugated anti-mouse IgG antibodies, and visualisation and quantification of spike protein labelling intensity by confocal microscopy. Plots show data expressed as mean ± SEM of three independent experiments. Solid line represents nonlinear regression using a 4-parameter logistic equation. Representative confocal images show labelling of cells under control conditions (in the absence of anti-spike antibody) or with spike protein pre-incubated with 10 μg/mL RBD1, RBD7 or RBD8.

In summary, we have developed a high content imaging assay to quantify SARS-CoV-2 spike protein binding to ACE2-expressing target cells. ACE2 expression in A549 cells was sufficient to detect various recombinant spike proteins from the Wuhan strain and from variants of concern Alpha, Delta and Omicron. The serine protease TMPRSS2 primes SARS-CoV-2 S protein for ACE2-dependent entry (5), but Omicron variants enter by a TMPRSS2-independent pathway (13). In our assay, TMPRSS2 expression did not offer an advantage to the detection of spike binding for spike proteins from TMPRSS2 entry-dependent variants.

We demonstrate that some commercially available monoclonal antibodies were able to neutralise recombinant spike binding in a manner comparable to replicating virus neutralisation.

Characterisation of spike (RBD)-specific antibodies is important to monitor immunological memory (14), and there is evidence that repertoires of SARS-CoV-2 epitopes targeted by antibodies vary according to severity of COVID-19 (15). While ELISA-based assays are important to characterise antibody repertoires by assessing recognition of immobilised spike proteins, the spike binding bioassay offers the advantage of a functional assay that assesses the ability to neutralise receptor binding without the need for high containment facilities.

As this assay system uses a ‘His-tag’ for detection, which is frequently used for purification on initial expression of a recombinant protein, His-tagged recombinant viral variant spike proteins are often rapidly available and accessible enabling the assay to be quickly adapted to test activity of therapeutics as new strains of SARS-CoV-2 emerge.

Across viral variants, although antibody affinity for spike protein may remain approximately constant, the affinity of spike protein for ACE2 may increase significantly and be associated with reduced antibody neutralisation (16). Measuring functional antibody blockade of the spike protein-ACE2 interaction is therefore likely to become increasingly important and we envisage that this cell-based, spike protein assay would complement existing biochemical and pseudovirus assays in the development of therapeutics, and may be adapted to other cell types to investigate potentially ACE2-independent binding of spike protein.

## Methods

### Antibodies and recombinant spike proteins

See Table 1 for a list of recombinant spike proteins used in this study. Hoechst-33342 was from ThermoFisher Scientific (H21492), His tag® antibody [HIS.H8] was from Abcam (ab18184), and goat anti-Mouse IgG H&L (Alexa Fluor® 488) was from ThermoFisher Scientific (A-11001). See table 2 for a list of antibodies against Spike protein.

### Cell culture

A549 lung carcinoma cells or A549 cells expressing human ACE2 or ACE2 and TMPRSS2 were obtained from Invivogen (a549, a549-hace2 or a549-hace2tpsa, respectively). Cells were cultured in F-12K Ham Nutrient Mixture media (ThermoFisher Scientific; 21127022) supplemented with 10% fetal bovine serum (Sigma-Aldrich; F9665) and 1% penicillin-streptomycin (ThermoFisher Scientific; 15140122) in a tissue culture incubator at 37 °C (5% CO_2_) using standard cell culture practices.

### Spike binding and blocking assays

Cells were seeded in 96-well plates overnight prior to incubation with recombinant spike proteins. Recombinant spike proteins, anti-His detection antibody, blocking antibodies or plasma were pre-incubated for 1 h prior to addition to cells for 1 h at 37 °C followed by preparation for confocal microscopy.

### Confocal Microscopy

Media was removed from cells and wells washed with PBS (ThermoFisher Scientific; 14190144). Cells were then fixed with 4% paraformaldehyde in PBS, permeabilized (ThermoFisher Scientific; 00-8333-56) prior to blocking (1% goat serum in PBS). Cells were then incubated with Alexa Fluor 488-conjugated anti-mouse IgG (1:500) and Hoechst (1:1000) before washing and imaging in PBS. Confocal microscopy was carried out using a Yokogawa CQ1 spinning-disc microscope using a x40 objective and appropriate excitation/emission settings for Hoechst and Alexa Fluor 488. Z-stack images were acquired and displayed as maximum intensity projections. Image analysis was carried out on the Yokogawa image analysis software.

### IgG/A/M ELISA

Levels of IgG, IgA and IgM antibodies in plasma samples were measured by protein ELISA using the IgG/A/M Sars-COV-2 ELISA kit following manufactures instructions (Binding Site; MK654).

### SARS-CoV-2 infection studies

Vero cells were cultured in DMEM supplemented with 10% foetal bovine serum, 2 mM l-glutamine, 100 U/mL penicillin, 10 μg/mL penicillin and 10 μg/mL streptomycin and 1% non-essential amino acids (cDMEM) and seeded into 96-well plates. SARS-CoV-2-England 2 (Wuhan strain) virus at 10^6^ IU/mL (GSAID Accession ID EPI_ISL_407073) was a kind gift from Christine Bruce, Public Health England. Antibodies were pre-incubated with virus for 1 h prior to addition to Vero cells. After a 48 h incubation at 37 °C cells were fixed with ice-cold methanol (5 min), washed with PBS and stained with rabbit anti-SARS-CoV-2 spike protein, subunit 1 (CR3022, The Native Antigen Company), detected by Alexa Fluor 555-conjugated goat anti-rabbit IgG secondary antibody (Invitrogen, ThermoFisher Scientific). Cell nuclei were stained with Hoechst 33342 (ThermoFisher Scientific). Cells were washed with PBS and then imaged and analysed using a ThermoFisher Scientific CellInsight CX5 High-Content Screening (HCS) platform. Infected cells and cell viability were detected by measuring perinuclear fluorescence above a set threshold determined by positive (untreated) and negative (uninfected) controls. Automated quantification algorithms were developed with assistance from Dr Henri Huppert, ThermoFisher Scientific, UK.

### Statistical analysis

Data are presented as mean + SEM or mean ± SEM from at least three independent experimental setups unless otherwise indicated. Curve-fitting data analysis was performed with KaleidaGraph (version 3.5).

### Ethics

All samples were obtained with informed consent and with approval from the appropriate Research Ethics Committee (REC Reference 20/WA/0216).

## Acknowledgements

This work was supported by Innovate UK [84361] and Celentyx Ltd. HJH and ZS are funded by a Medical Research Foundation intermediate career fellowship to ZS (UKRI, Grant number MRF-169-0001-F-STAM-C0826).

## Author Contributions

Neale Harrison (conceptualization, methodology, investigation, resources, writing, formal analysis, visualization, supervision), Lauren Richardson (methodology, investigation, writing, formal analysis, visualization), Chiara Pallini (methodology, investigation, writing), Ines Morano (methodology, investigation, writing), Elizabeth Jinks (methodology, investigation, writing), Jamie Cowley (methodology, investigation, writing), Hujo Chan (methodology, investigation, writing, formal analysis), Harriet J Hill (methodology, investigation, formal analysis, writing), Cristina Matas de las Heras (methodology, investigation, writing), Ana Teodosio (methodology, investigation, writing), Andrea S Lavado (methodology, investigation, writing), Timothy R Dafforn (conceptualization, methodology, writing, funding acquisition), Dimitris K Grammatopoulos (conceptualization, methodology, writing, funding acquisition), John Gordon (conceptualization, methodology, writing, funding acquisition), Catherine A Brady (conceptualization, methodology, writing, funding acquisition), Lawrence S Young (conceptualization, methodology, writing, funding acquisition), Nicholas M Barnes (conceptualization, methodology, writing, funding acquisition), Zania Stamataki (conceptualization, methodology, writing, funding acquisition) and Omar S Qureshi (conceptualization, methodology, resources, writing, formal analysis, visualization, supervision).

## References

1. Robinson, P. C., Liew, D. F. L., Tanner, H. L., Grainger, J. R., Dwek, R. A., Reisler, R. B., Steinman, L., Feldmann, M., Ho, L.-P., Hussell, T., Moss, P., Richards, D., and Zitzmann, N. (2022) COVID-19 therapeutics: Challenges and directions for the future. Proceedings of the National Academy of Sciences. 10.1073/pnas.2119893119

2. Hwang, Y.-C., Lu, R.-M., Su, S.-C., Chiang, P.-Y., Ko, S.-H., Ke, F.-Y., Liang, K.-H., Hsieh, T.-Y., and Wu, H.-C. (2022) Monoclonal antibodies for COVID-19 therapy and SARS-CoV-2 detection. Journal of Biomedical Science. 10.1186/s12929-021-00784-w

3. Galipeau, Y., Greig, M., Liu, G., Driedger, M., and Langlois, M.-A. (2020) Humoral Responses and Serological Assays in SARS-CoV-2 Infections. Frontiers in Immunology. 10.3389/fimmu.2020.610688

4. Ramesh, S., Govindarajulu, M., Parise, R. S., Neel, L., Shankar, T., Patel, S., Lowery, P., Smith, F., Dhanasekaran, M., and Moore, T. (2021) Emerging SARS-CoV-2 Variants: A Review of Its Mutations, Its Implications and Vaccine Efficacy. Vaccines. 9, 1195

5. Hoffmann, M., Kleine-Weber, H., Schroeder, S., Krüger, N., Herrler, T., Erichsen, S., Schiergens, T. S., Herrler, G., Wu, N.-H., Nitsche, A., Müller, M. A., Drosten, C., and Pöhlmann, S. (2020) SARS-CoV-2 Cell Entry Depends on ACE2 and TMPRSS2 and Is Blocked by a Clinically Proven Protease Inhibitor. Cell. 181, 271–280

6. Wang, B., Zhao, J., Liu, S., Feng, J., Luo, Y., He, X., Wang, Y., Ge, F., Wang, J., Ye, B., Huang, W., Bo, X., Wang, Y., and Jeff, J. (2022) ACE2 Decoy Receptor Generated by High-throughput Saturation Mutagenesis Efficiently Neutralizes SARS-CoV-2 and Its Prevalent Variants. Emerging Microbes & Infections. 10.1080/22221751.2022.2079426

7. Shah, M., Ung Moon, S., Hyun Kim, J., Thanh Thao, T., and Goo Woo, H. (2022) SARS-CoV-2 pan-variant inhibitory peptides deter S1-ACE2 interaction and neutralize delta and omicron pseudoviruses. Computational and Structural Biotechnology Journal. 20, 2042–2056

8. Zhang, L., Narayanan, K. K., Cooper, L., Chan, K. K., Devlin, C. A., Aguhob, A., Shirley, K., Rong, L., Rehman, J., Malik, A. B., and Procko, E. (2022) An engineered ACE2 decoy receptor can be administered by inhalation and potently targets the BA.1 and BA.2 omicron variants of SARS-CoV-2. 10.1101/2022.03.28.486075

9. Shah, M., and Woo, H. G. (2022) Omicron: A Heavily Mutated SARS-CoV-2 Variant Exhibits Stronger Binding to ACE2 and Potently Escapes Approved COVID-19 Therapeutic Antibodies. Frontiers in Immunology. 10.3389/fimmu.2021.830527

10. Walls, A. C., Park, Y.-J., Tortorici, M. A., Wall, A., McGuire, A. T., and Veesler, D. (2020) Structure, function, and antigenicity of the sars-cov-2 spike glycoprotein. Cell. 181, 281–292

11. Dejnirattisai, W., Huo, J., Zhou, D., Zahradník, J., Supasa, P., Liu, C., Duyvesteyn, H. M. E., Ginn, H. M., Mentzer, A. J., Tuekprakhon, A., Nutalai, R., Wang, B., Dijokaite, A., Khan, S., Avinoam, O., Bahar, M., Skelly, D., Adele, S., Johnson, S. A., and Amini, A. (2022) SARS-CoV-2 Omicron-B.1.1.529 leads to widespread escape from neutralizing antibody responses. Cell. 10.1016/j.cell.2021.12.046

12. Tandon, R., Mitra, D., Sharma, P., McCandless, M. G., Stray, S. J., Bates, J. T., and Marshall, G. D. (2020) Effective screening of SARS-CoV-2 neutralizing antibodies in patient serum using lentivirus particles pseudotyped with SARS-CoV-2 spike glycoprotein. Scientific Reports. 10.1038/s41598-020-76135-w

13. Meng, B., Abdullahi, A., Ferreira, I. A. T. M., Goonawardane, N., Saito, A., Kimura, I., Yamasoba, D., Gerber, P. P., Fatihi, S., Rathore, S., Zepeda, S. K., Papa, G., Kemp, S. A., Ikeda, T., Toyoda, M., Tan, T. S., Kuramochi, J., Mitsunaga, S., Ueno, T., and Shirakawa, K. (2022) Altered TMPRSS2 usage by SARS-CoV-2 Omicron impacts tropism and fusogenicity. Nature. 10.1038/s41586-022-04474-x

14. Dan, J. M., Mateus, J., Kato, Y., Hastie, K. M., Yu, E. D., Faliti, C. E., Grifoni, A., Ramirez, S. I., Haupt, S., Frazier, A., Nakao, C., Rayaprolu, V., Rawlings, S. A., Peters, B., Krammer, F., Simon, V., Saphire, E. O., Smith, D. M., Weiskopf, D., and Sette, A. (2021) Immunological Memory to SARS-CoV-2 Assessed for Up to 8 Months After Infection. Science. 10.1126/science.abf4063

15. Gregory, D. J., Vannier, A., Duey, A. H., Roady, T. J., Dzeng, R. K., Pavlovic, M. N., Chapin, M. H., Mukherjee, S., Wilmot, H., Chronos, N., Charles, R. C., Ryan, E. T., LaRocque, R. C., Miller, T. E., Garcia-Beltran, W. F., Thierauf, J. C., Iafrate, A. J., Mullenbrock, S., Stump, M. D., and Wetzel, R. K. (2022) Repertoires of SARS-CoV-2 epitopes targeted by antibodies vary according to severity of COVID-19. Virulence. 13, 890–902

16. Bachmann, M. F., Mohsen, M. O., and Speiser, D. E. (2022) Increased receptor affinity of SARS-CoV-2: a new immune escape mechanism. npj Vaccines. 10.1038/s41541-022-00479-9

